# Mechanistic evidence that motif-gated domain recognition drives contact prediction in protein language models

**DOI:** 10.1101/2025.08.22.671739

**Authors:** Jatin Nainani, Bryn Marie Reimer, Connor Watts, David Jensen, Anna G. Green

**Affiliations:** Manning College of Information and Computer Sciences, University of Massachusetts, Amherst; Queen Mary University of London

## Abstract

Protein language models (pLMs) achieve state-of-the-art performance on protein structure and function prediction tasks, yet their internal computations remain opaque. Sparse autoencoders (SAEs) have been used to recover sparse features, called latents, from pLM layer representations, whose activations correlate with known biological concepts. However, prior work has not established which model concepts are *causally necessary* for pLM performance on down-stream tasks. Here, we adapt causal activation patching to the pLM setting and perform it in SAE latent space to extract the minimal circuit responsible for accuracy in a contact prediction task for two case study proteins. We observe that preserving only a tiny fraction of latent–token pairs (0.022% and 0.015%) is sufficient to retain contact prediction accuracy in a residue unmasking experiment. Our circuit indicates a two-step computation in which early-layer *motif detectors* respond to short local sequence patterns, gating mid-to-late *domain detectors* which are selective for protein domains and families. Path-level ablations confirm the causal dependence of domain latents on upstream motif latents. To evaluate these components quantitatively, we introduce two diagnostics: a *Motif Conservation Test* and a *Domain Selectivity Framework* that supports hypothesis-driven tests. All candidate motif-detector latents pass the conservation test, and 18/23 candidate domain-detector latents achieve AUROC ≥0.95. To our knowledge, this is the first circuits-style causal analysis for pLMs, pin-pointing the motifs, domains, and motif-domain interactions that drive contact prediction in two specific case studies. The framework introduced herein will enable future mechanistic dissection of protein language models. Code available at https://github.com/NainaniJatinZ/plm_circuits

## 1 Introduction

Protein language models (pLMs) now sit at the core of modern computational biology, achieving strong performance at many computational biology tasks [Lin et al., 2023a, Wu et al., 2022, Elnaggar et al., 2022, Nijkamp et al., 2023, Ullanat et al., 2025]. Yet, we know little about the computational mechanisms that enable these networks to transform raw amino acid sequences into higher-level structural or functional inferences. Without insight into a model’s internal computations, we cannot effectively reason about predictions, debug systematic errors, or extract new biological insights.

The field of mechanistic interpretability provides insight into model computations by reverse-engineering neural networks at feature-level resolution. A central obstacle is that most neu-rons in large language models are polysemantic: a single neuron fires for several unrelated sequence features [Olah et al., 2020, Elhage et al., 2022], which hampers interpretation. *Sparse autoencoders* (SAEs) project the dense activations of neurons in a large language model into a higher-dimensional sparse space, where each neuron—called a latent—is intended to fire selectively for a single concept [Bricken et al., 2023, Gao et al., 2024, Templeton, 2024]. Efforts to port SAEs to biological sequence models have provided descriptive evidence that pLMs contain features for sequence motifs, secondary-structure elements, and whole protein families [Simon and Zou, 2024, Adams et al., 2025, Gujral et al., 2025]. While SAEs reveal what concepts a model encodes, only causal interventions— perturbing a latent’s activation and measuring the effect on the output—can tell us whether a model depends on that concept (as encoded by a specific latent) to make predictions [Vig et al., 2020].

Here, we extend the causal interpretability framework [Lindsey et al., 2025] to SAE latents in pLMs. We study the residue–residue contact prediction capabilities of ESM-2, using causal activation patching to measure the contribution of each interpretable SAE latent. We show that contact prediction depends on only a small subset of representations by identifying specific latents, active at specific token positions, that are individually necessary. We provide evidence that the causally necessary representations correspond to biologically meaningful concepts. We show that early layers detect short sequence motifs that causally gate domain recognition in deeper layers, forming a multi-step computational circuit. By tracing ESM-2’s contact prediction for specific proteins back to a compact and interpretable subgraph of the model, and by publishing the tools needed to extend this analysis, we move towards mechanistic understanding of protein language models.

## 2 Results

### 2.1 A framework for causal circuit discovery in pLMs

Here we outline a framework for causal circuit discovery in pLMs, building on established interpretability practice in language models [Wang et al., 2022, Lindsey et al., 2025]. Our analysis follows four steps: (i) represent hidden activations in an interpretable sparse latent space, (ii) define a task for the pLM of interest to perform, (iii) perform causal interventions in the interpretable latent space to perturb performance of the task, and (iv) interpret the biological meaning, if any, of the observed causal latents. We briefly describe each choice below.

#### (i) Sparse autoencoders provide an interpretable latent space for pLMs

Recent studies [Adams et al., 2025, Simon and Zou, 2024, Gujral et al., 2025] have shown that sparse autoencoders (SAEs) [Elhage et al., 2022] project the dense, polysemantic activations of protein language models into interpretable sparse representations where individual neurons, called latents, capture meaningful biological concepts. We use SAEs from Adams et al.: eight layers (4–32 in steps of 4), 4,096 latents each.

#### (ii) Contact prediction as an example task for PLMs

An ideal downstream task for activation patching to interpret pLM behavior is one where the model exhibits a near-discrete behavior switch over a small change in input, because this minimizes possible confounds[Zhang and Nanda, 2023, Wang et al., 2022]. Such behavior has been documented for the residue-residue contact prediction capabilities of ESM-2-3B. Zhang et al. showed that contact recovery for distant secondary-structure elements (SSEs) in a partially masked sequence stays near random until a critical number of residues flanking the SSEs are unmasked; then accuracy “jumps” to near-perfect (Fig. 1A-C). This results in two nearly identical inputs for which the network has a major transition in its output. We select two such case-study proteins: MetXA (UniProt: P45131), where unmasking two additional residues raises the contact recovery score *m*(*X*) from 0.02 to 0.58; and TOP2 (UniProt: P06786), where unmasking four extra residues raises *m*(*X*) from 0.06 to 0.86.

**Figure 1.**
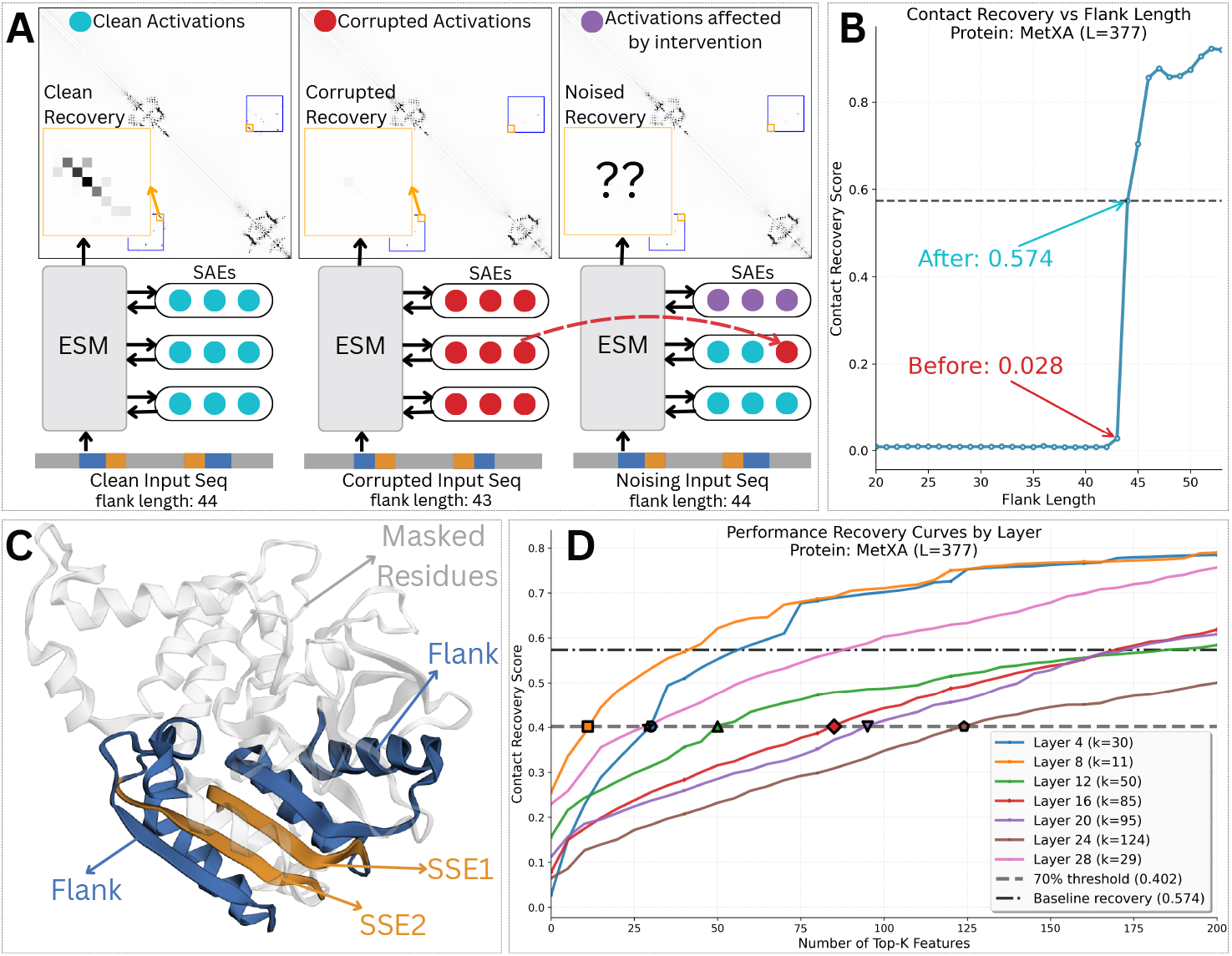
**(A)** Residue-residue contact maps for MetXA under three conditions: clean sequence (44 aa) yields near-perfect recovery; corrupted sequence (43 aa) yields near-random contact recovery; causal intervention replaces clean activation with a single latent patched to its corrupted value. Loss of contact recovery flags that latent as causal. **(B)** Contact recovery jump behavior reproduced from Zhang et al. [2024]. **(C)** MetXA structure (RCSB-PDB ID: 2B61 Chain A [Mirza et al., 2005]); the studied SSE elements are highlighted in yellow. **(D)** Recovery vs. per-layer circuit size. Starting points differ by error-node performance *m*_0_.

#### (iii) Activation patching to identify causal network components

Activation patching [Vig et al., 2020, Finlayson et al., 2021] can measure the causal necessity of every latent, at each token position (which we call “latent-token pair”) for a specific task and input pair. For each case study input pair, we define the *corrupted input* as the input sequence and with low contact recovery, and the *clean input* as the related sequence and with high recovery. The corresponding latent activations are also called corrupted and clean, respectively. We use activation patching to compute the Indirect Effect (IE), which measures the causal influence of the input on the output through a single latent acting as a mediator Pearl [2022], by patching corrupted activations into a clean pass of the model to determine whether specific latent-token pairs were causally necessary for the task (Methods, Sec. 4; Fig. 1A). We identify a “circuit”: the smallest set of latents whose clean activations sustain contact recovery (Sec. 2.2). We later extend this analysis to interactions between components (Sec. 2.4.3).

#### (iv) Interpretation of causal network components

After identification of relevant latent-token pairs, we ask what biological computations they represent and how they interact to produce the jump in conctact recovery. We proceed in two stages. First, we assign *global* semantic labels by inspecting each latent’s top-activating residues and proteins across the corpus. Second, we characterize each latent’s *task-specific role* by comparing its activation in the clean vs. corrupted inputs around the flanking regions that control the jump. From these observations we formulate hypotheses about (i) the function of individual latents and (ii) directed interactions between latents across layers (Sec. 2.3). We then attempt to falsify these hypotheses using targeted interventions and selectivity tests (Sec. 2.4).

### 2.2 Causal circuits underlying contact prediction for two case study proteins

We compute the indirect effect (IE) for 4,096 latents across 8 layers at *∼*400 token positions (≈ 1.3 *×* 10^7^ latent–token pairs), and then compute both global and layer-wise circuits.

#### Global circuits for quantification

We ask for the smallest subset *K* of latent–token pairs whose clean activations keep contact recovery above a threshold *θ* of the clean score (Fig. 1D). For each protein, we fix the threshold at 70% of the clean contact recovery, *θ* = 0.70 × *m*_clean_. Then, for a given *K*, we patch all non–top *K* pairs to their corrupted activations, recompute the model, and check whether *m*(*X*) ≥ *θ*; if so, those *K* latent-token pairs are sufficient (Sec. 4). Fig. 13a and Fig. 14a show the increase in *m*(*X*) with increasing top *K* pairs considered. For both proteins, only a tiny subgraph is needed for the circuit to reach the 70%-of-clean threshold *θ*: 2,401 pairs (0.022%) for MetXA and 1,801 (0.015%) for TOP2. Thus the contact-prediction switch is governed by a subgraph three orders of magnitude smaller than the full network.

#### Layer-wise circuits for interpretability

Manually inspecting thousands of pairs is infeasible without a prior, so we analyze one layer at a time. For layer *ℓ*, the bottleneck ℬ_*ℓ*_ is the smallest within-layer subset that maintains *m*(*X*) ≥ *θ* with all other layers left unmodified. Because the SAEs cannot perfectly reconstruct activations, SAEs include an “error” node that carries reconstruction loss [Marks et al., 2024]. For layer *ℓ*, let the zero-circuit performance *m*_0_(*ℓ*) be the score when all pairs in that layer are patched to their corrupted values and only the error node remains active. Because this node contains the activations “unexplained” by the SAE, we treat it as non-interpretable; continuing work aims to reduce its contribution [Rajamanoharan et al., 2024, Bussmann et al., 2024]. We focus on the *explainable window W*_*ℓ*_ = *θ ™ m*_0_(*ℓ*), the margin above the error-only baseline that a layer’s bottleneck must account for; we report drops both in absolute units and as a percent of *W*_*ℓ*_. The resulting bottlenecks ℬ_*ℓ*_ contains on average only ≈60 latent–token pairs—tractable for qualitative study yet still drawn from the very top of the global IE ranking (Fig. 1D).

### 2.3 Manual inspection of two case studies reveals a motif-gated, domain-recognition circuit

We now manually annotate the causally relevant latents in each layer-wise bottleneck for both of our case study proteins. Manual annotation in this section will be used to generate hypotheses that are quantitatively tested in the next section. For every latent–token pair in each layer’s bottleneck we ask two questions. First, (Q1) what biological signal, if any, does the latent usually represent? Second, (Q2) how does that latent’s activation change from the corrupted to the clean input, and why might that change unblock the downstream domain detector? We answer both questions by (E1) inspecting the 20 UniRef50 proteins that most strongly activate the latent and (E2) comparing the latent’s activation maps between corrupted and clean runs of the case-study protein (Sec. 4). We describe motifs using the following notation: specific residues use one-letter aminoacid codes; *X* denotes any residue; an <u>underline</u> marks the token where the latent activates (e.g., XXI indicates a latent that activates two residues upstream of an isoleucine). See Sec. B.3 for details.

#### 2.3.1 Homoserine *O*-acetyltransferase

Unmasking I133 (left) and F363 (right) raises contact recovery from *m*_corr_ = 0.02 to *m*_clean_ = 0.58 (Δ*m* = 0.56); thus the threshold *T* = 0.40. We provide per-latent and per-latent-cluster details in Table 1 and Table 3 respectively. Latents not detailed could not readily be assigned a global or task-specific role by manual inspection.

**Table 1.** Layer-wise role for latents in MetXA circuit.

| Layer | Role | Class | Quantity | % of Layer | $\Delta m$ drop (abs) | $\Delta m$ drop (rel % w/o err) |
| --- | --- | --- | --- | --- | --- | --- |
| 4<br>(Zero Circuit Base-line: 0.027) | Motif Detectors | Direct | 5 | 16.7 | 0.0860 | 22.9 |
|  |  | Indirect | 5 | 16.7 | 0.0500 | 13.3 |
|  | Total Explained | - | 10 | 33.3 | 0.1330 | 35.3 |
|  | Unlabeled | - | 20 | 66.6 | - | - |
| 8<br>(Zero Circuit Base-line: 0.255) | Domain Detectors | Annotated | 1 | 9.1 | 0.0150 | 9.5 |
|  |  | Misc | 4 | 36.7 | 0.0590 | 38.0 |
|  | Total Explained | - | 5 | 45.4 | 0.0740 | 47.2 |
|  | Unlabeled | - | 6 | 54.6 | - | - |
| 12<br>(Zero Circuit Base-line: 0.156) | Domain Detectors | Annotated | 2 | 4.0 | 0.0190 | 7.7 |
|  |  | Misc | 6 | 12.0 | 0.0720 | 29.1 |
|  | Total Explained | - | 8 | 16.0 | 0.0930 | 37.7 |
|  | Unlabeled | - | 42 | 84.0 | - | - |

##### Layer 4

The zero-circuit performance *m*_0_ = 0.027, thus the explainable window *W* = *T ™ m*_0_ = 0.373. The circuit requires 30 latent–token pairs to meet the criterion. We find latents that activate on short sequence patterns (Q1) across the proteome (E1), so we label them *motif detectors*. This cluster contains 10 pairs (33.33% of the layer). We identify *direct* motif detectors that include one of the two newly unmasked residues (16.67% of layer). Ablating them reduces *m*(*X*) by 22.9% of *W* (0.0863). For example, a latent at P131 fires on the motif XXI across its top 20 activating proteins (E1, Fig. 2D) and switches on only when I133 is revealed (E2→Q2, Fig. 2C). We also identify *indirect* motif detectors, which activate on residues not in the flank region, but whose activation differs between corrupted and clean inputs (16.67% of layer). Ablating them reduces *m*(*X*) by 13.3% of *W* (0.0501). Example: a latent at I181 (center of SSE1) activates for PXXXXXX (E1); its activation rises once distant flanks are unmasked (E2→Q2). Together, motif detectors account for 35% of *W* (0.132).

**Figure 2.**
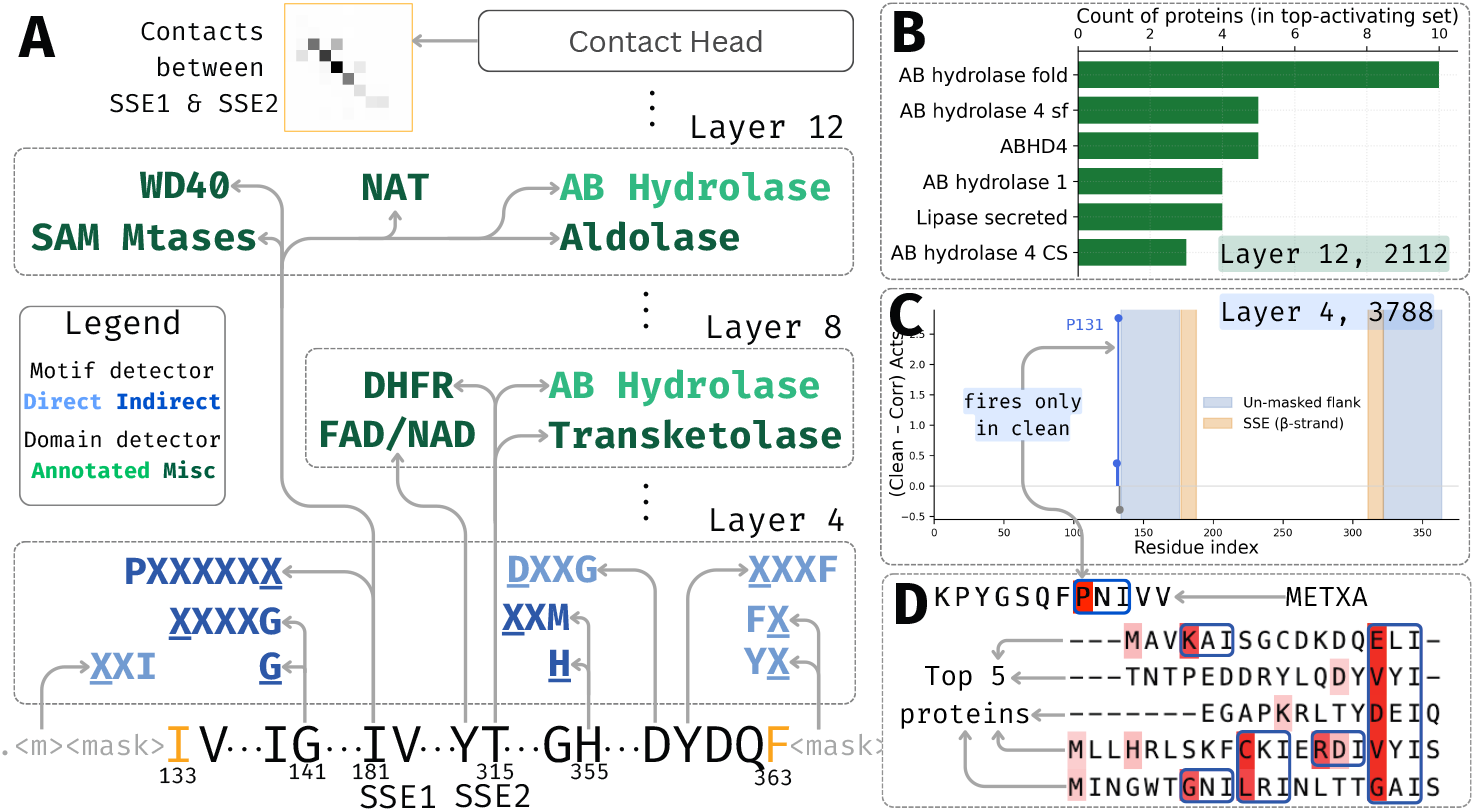
**(A)** Motif and domain detectors in the layer-wise circuit for MetXA. Arrows connect causal latent-token pairs. **(B)** Frequency of Interpro domains for top 10 proteins with highest activation for latent 2112 in layer 12. **(C)** Change in activation of latent 3788 in layer 4 from clean to corrupted. The latent-token pair at position P131 only activates when I133 is unmasked in the clean sequence supporting motif activation of XXI. **(D)** Screenshot of activation patterns of the top 5 proteins for latent 3788 in layer 4 from Interprot [Adams et al., 2025].

##### Layer 8

The zero-circuit performance is *m*_0_ = 0.255, thus the explainable window *W* = *T ™ m*_0_ = 0.145. The circuit requires 11 latent–token pairs to meet the criterion. We observe latents which which activate on proteins (Q1) containing a specific domain annotation, and term them *domain detectors*. This cluster contains 5 pairs (45.45% of the layer). Domains matching MetXA ‘s own annotation (*annotated* domain detectors) comprise 9.09% of layer, and ablating them reduces *m*(*X*) by 9.54% of *W* (0.015). We also see latents that activate on other domains (*miscellaneous* domain detectorrs), such as DHFR, FAD/NAD, or transketolase domains (E1) (36.36% of layer). Ablating them reduces *m*(*X*) by 38% of *W* (0.0597), suggesting the model cross-checks related folds. Together, domain detectors account for 47.2% of *W* (0.0742).

##### Layer 12

The zero-circuit performance is *m*_0_ = 0.156, thus the explainable window *W* = *T ™ m*_0_ = 0.244. The circuit requires 50 latent–token pairs to meet the criterion. Similar to Layer 8, Layer 12 contains *domain detectors*. The group of domain detectors comprises 8 pairs (16% of the layer). *Annotated* domain detectors comprise 4% of the layer, and ablating them reduces *m*(*X*) by 7.74% of *W* (0.0192). For example, a latent selective for the AB-hydrolase fold (E1, Fig. 2B) is causal at two token positions. *Miscellaneous* domain detectors activate on NAT, SAM-methyltransferase, WD40, and aldolase families (E1) (12% of layer). Ablating them reduces *m*(*X*) by 29.11% of *W* (0.072). Together, domain detectors account for 37.7% of *W* (0.093).

#### 2.3.2 DNA topoisomerase 2

Unmasking T31–Y32 (left) and G271–E272 (right) raises contact recovery from *m*_corr_ = 0.06 to *m*_clean_ = 0.86 (Δ*m* = 0.80); threshold *T* = 0.6. Per-cluster and per-latent details are in Table 2 and Table 4. SAE error nodes already achieve *m*_0_ = 0.59 at Layer 8 (*> T*), so we analyze Layers 4, 12, and 16 where *W >* 0. As in MetXA, Layer 4 contains motif detectors and Layers 12/16 contain domain detectors. The novelty is not the presence of these two functional classes but their reappearance with protein-appropriate content: TOP2’s Layer 4 motifs differ from MetXA’s, and its domain detectors align with TOP2’s GHKL/HATPase_c annotation. We cover the per-layer analysis in the Appendix A.

**Table 2.**
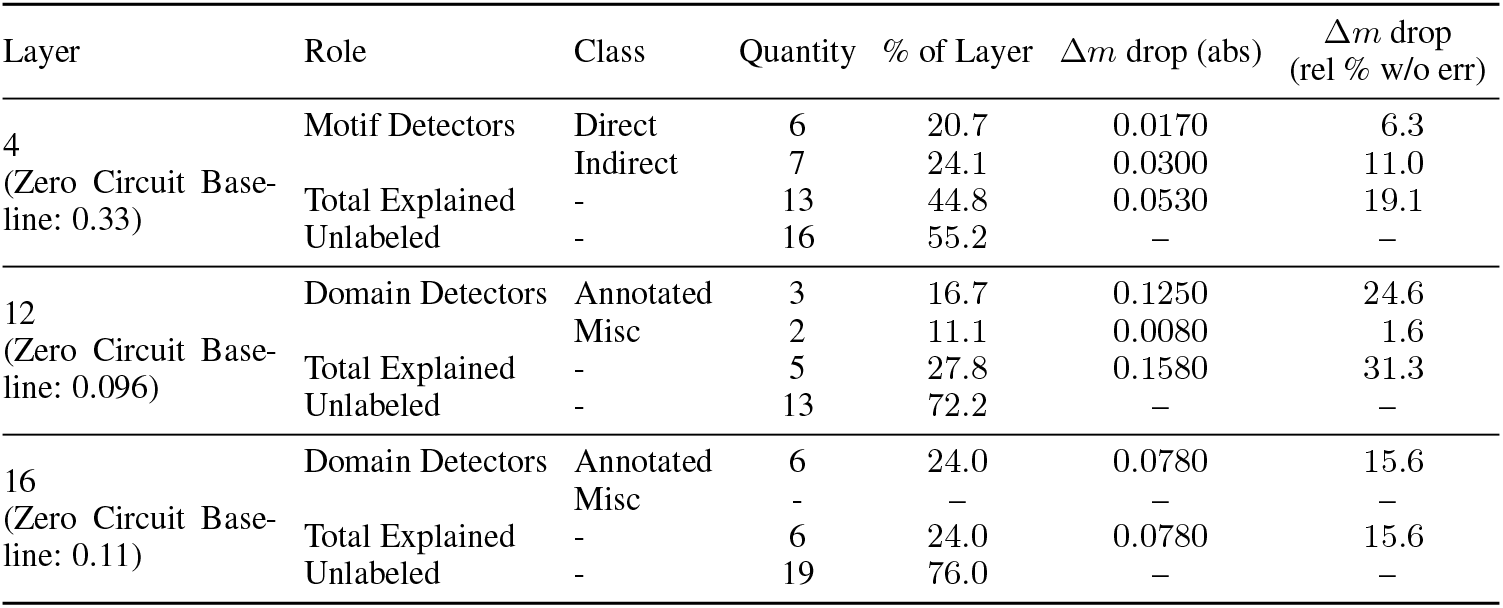
Layer-wise role for latents in TOP2 circuit.

**Table 3.**
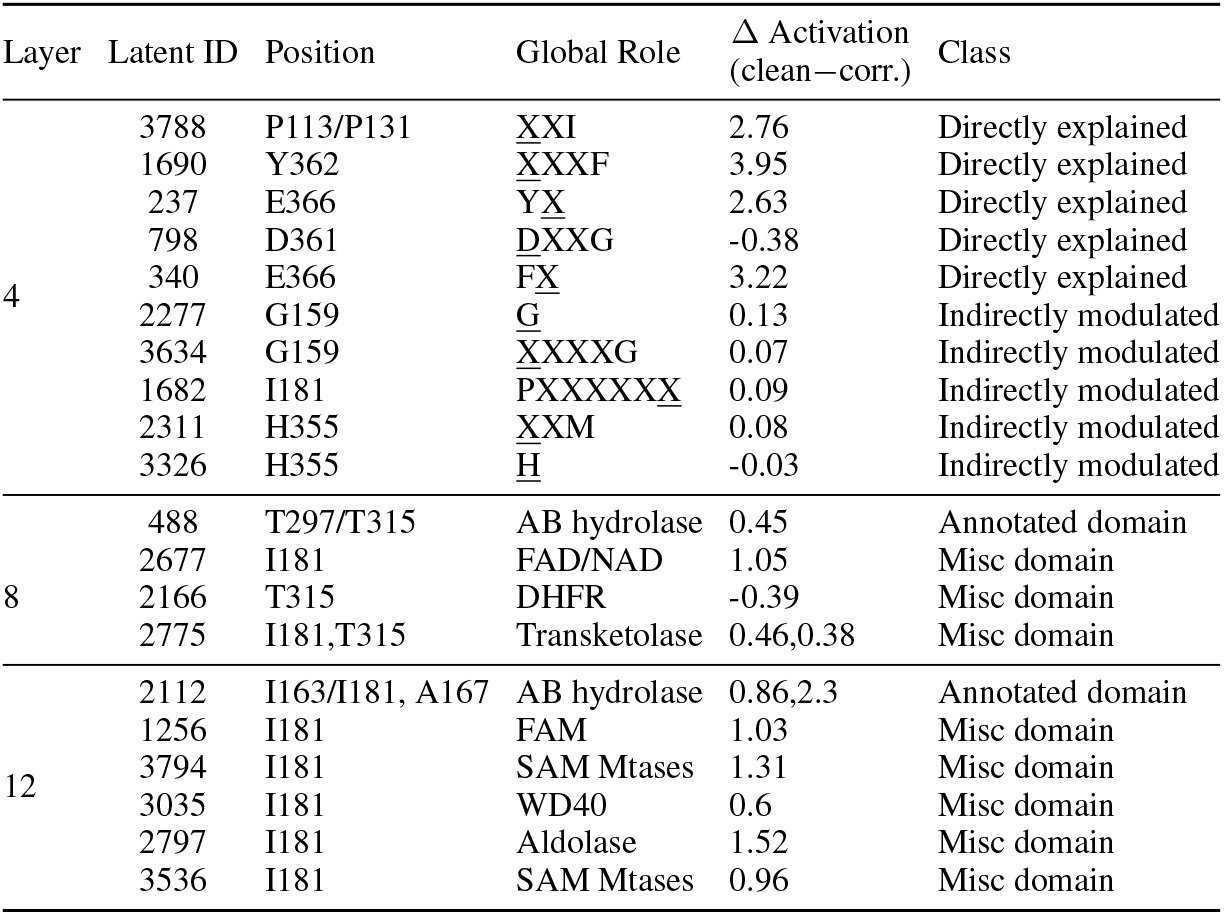
Latent census by layer (MetXA).

**Table 4.**
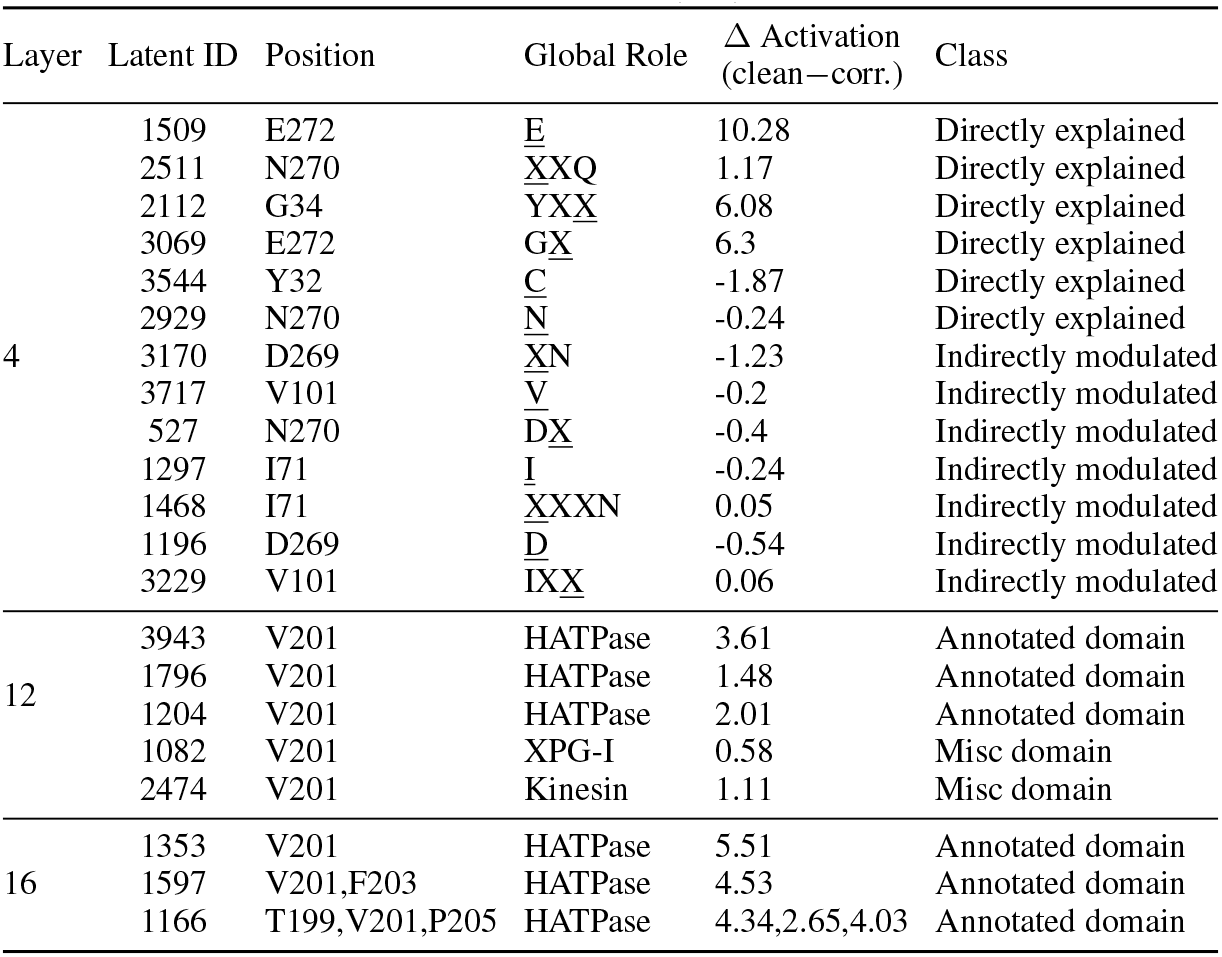
Latent census by layer (TOP2).

**Table 5.**
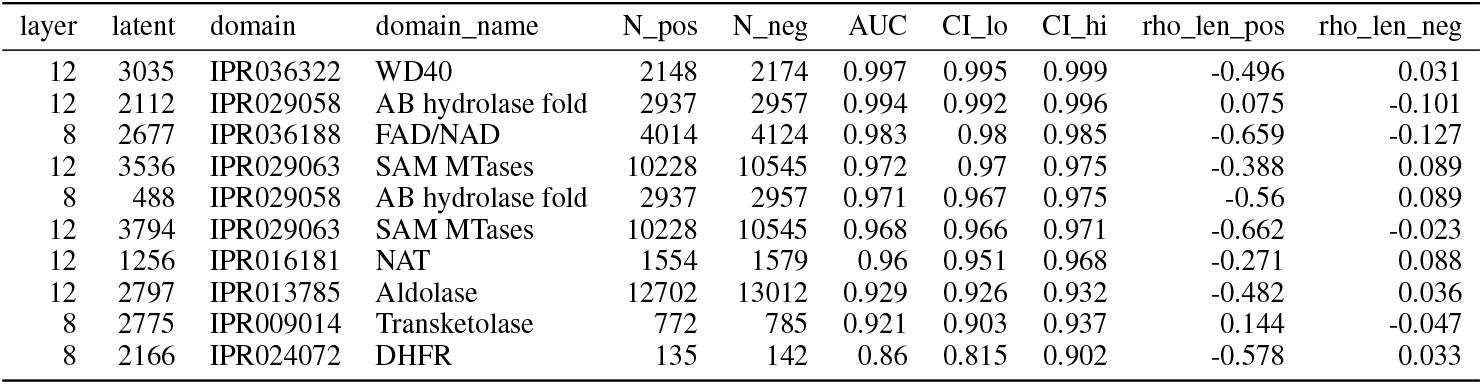
Top domain selectivity latents by AUC for aggregator top-*q*, group MetXA. We show the number of positive and negative samples, bootstrap 95% CI, spearmann rho correlation of the score vs length of sample.

**Table 6.** Top domain selectivity latents by AUC for aggregator top-*q*, group TOP2. We show the number of positive and negative samples, bootstrap 95% CI, spearmann rho correlation of the score vs length of sample.

| layer | latent | domain | domain_name | N_pos | N_neg | AUC | CI_lo | CI_hi | rho_len_pos | rho_len_neg |
| --- | --- | --- | --- | --- | --- | --- | --- | --- | --- | --- |
| 12 | 2472 | IPR036961 | Kinesin | 221 | 222 | 0.998 | 0.992 | 1 | -0.292 | -0.028 |
| 16 | 3077 | IPR036890 | HATPase_C_sf | 1700 | 1737 | 0.997 | 0.995 | 0.999 | -0.53 | 0.011 |
| 16 | 1353 | IPR036890 | HATPase_C_sf | 1700 | 1737 | 0.996 | 0.994 | 0.998 | -0.606 | 0.013 |
| 16 | 1814 | IPR036890 | HATPase_C_sf | 1700 | 1737 | 0.996 | 0.993 | 0.998 | -0.206 | 0.091 |
| 12 | 1145 | IPR036890 | HATPase_C_sf | 1700 | 1737 | 0.996 | 0.993 | 0.998 | -0.356 | -0.047 |
| 20 | 2311 | PF13589 | HATPase_c_3 | 869 | 892 | 0.995 | 0.992 | 0.999 | 0.076 | 0.048 |
| 12 | 3943 | IPR036890 | HATPase_C_sf | 1700 | 1737 | 0.995 | 0.993 | 0.997 | -0.588 | 0.058 |
| 12 | 1796 | IPR036890 | HATPase_C_sf | 1700 | 1737 | 0.994 | 0.991 | 0.997 | -0.107 | 0.115 |
| 12 | 1082 | PF00867 | XPG_I | 220 | 223 | 0.994 | 0.983 | 1 | -0.062 | -0.055 |
| 16 | 1166 | PF13589 | HATPase_c_3 | 869 | 892 | 0.992 | 0.987 | 0.997 | 0.135 | 0.015 |
| 12 | 1204 | IPR036890 | HATPase_C_sf | 1700 | 1737 | 0.974 | 0.969 | 0.978 | 0.409 | 0.094 |
| 16 | 1597 | IPR036890 | HATPase_C_sf | 1700 | 1737 | 0.794 | 0.778 | 0.811 | 0.487 | 0.077 |
| 16 | 3994 | PF13589 | HATPase_c_3 | 869 | 892 | 0.712 | 0.685 | 0.739 | 0.124 | 0.007 |

#### 2.3.3 Shared mechanism and hypotheses

Across both proteins, we observe that early layers (L4) contain motif detectors; mid/late layers (L8-16) contain domain detectors. Unmasking flank residues raises activation in both *direct* and *indirect* motif detectors, and domain detectors. The reason for activation of indirect motif detectors and domain detectors is not readily obvious, but one hypothesis is that the motif detectors give signal to the domain detectors. Because the explainable windows reported here are layer-specific and not cumulative, we test end-to-end causal gating with path-level ablations in Sec. 2.4.3.

Together, the case studies suggest a two-step circuit, that we hypothesize works in the following way: (1) early-layers require latents that detect specific motifs, (2) later layers require latents that detect specific domains, and (3) domain recognition is causally gated by the early layer motif detectors.

### 2.4 Quantitative validation of mechanistic hypotheses

In this section, we subject our hypotheses from the case studies to quantitative validation.

#### 2.4.1 Motif detectors preferentially activate on assigned motifs

Our case studies suggested that Layer 4 latents function as specific motif detectors which trigger the contact prediction circuit. We now ask whether each latent does indeed fire on a specific motif across all proteins, not just for our specific case study proteins.

We sample 10,000 proteins from the set of UniProt reviewed proteins and record the activation for each latent for each input token (Sec. 4.2.1). For every latent in our layer-specific bottlenecks, we create a sequence logo by recording the window around top activating token for each protein. For each latent in our qualitative analysis that was seen to be associated with a particular motif, that motif was also present in the sequence logo and accounted for more than 50% of the information at each position. As shown in Fig 3, we see that the residue that is fixed for a motif (*e*.*g*. F in XXXF) and its position are highly conserved whereas the rest of the window is not. All sequence logos are displayed in Fig. 5 and 6.

**Figure 3.**
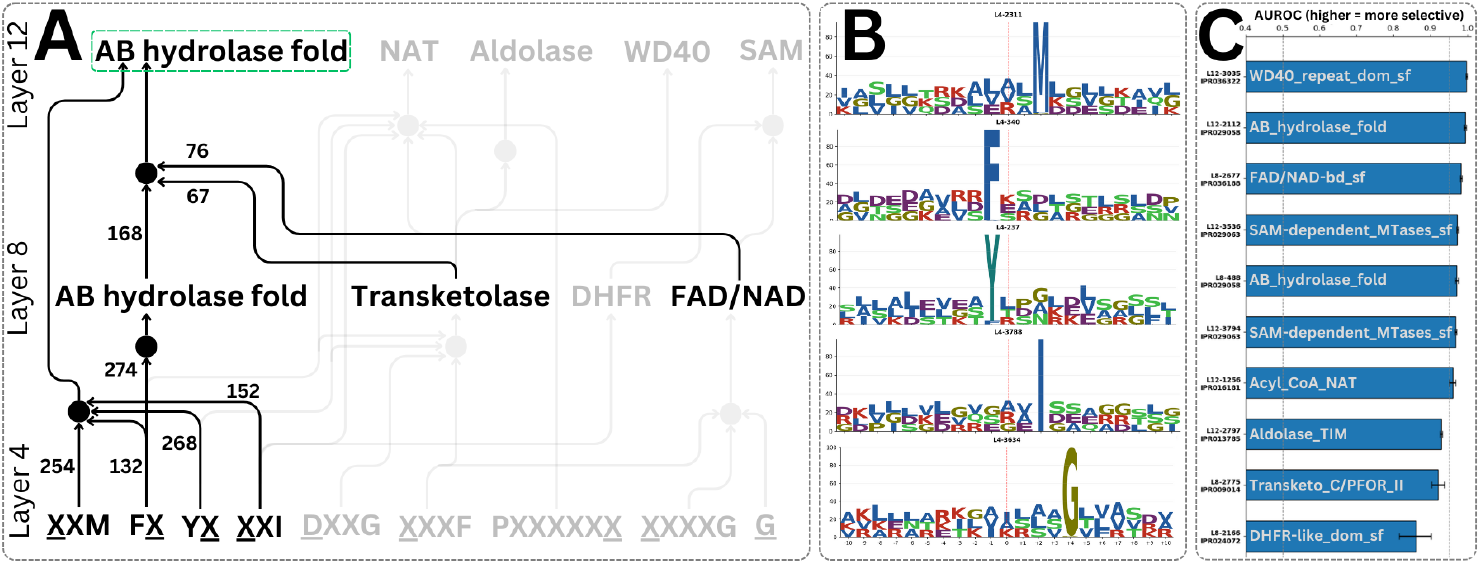
**(A)** Circuit diagram for MetXA with causal edges between latents. The subgraph for AB hydrolase is highlighted for readability, with edge ranks (out of 1580) shown. **(B)** Sequence logos for hypothesized motif detectors. **(C)** AUROC scores for hypothesized domain detectors. Label on the bar denotes the “short name” on Interpro.

**Figure 4.**
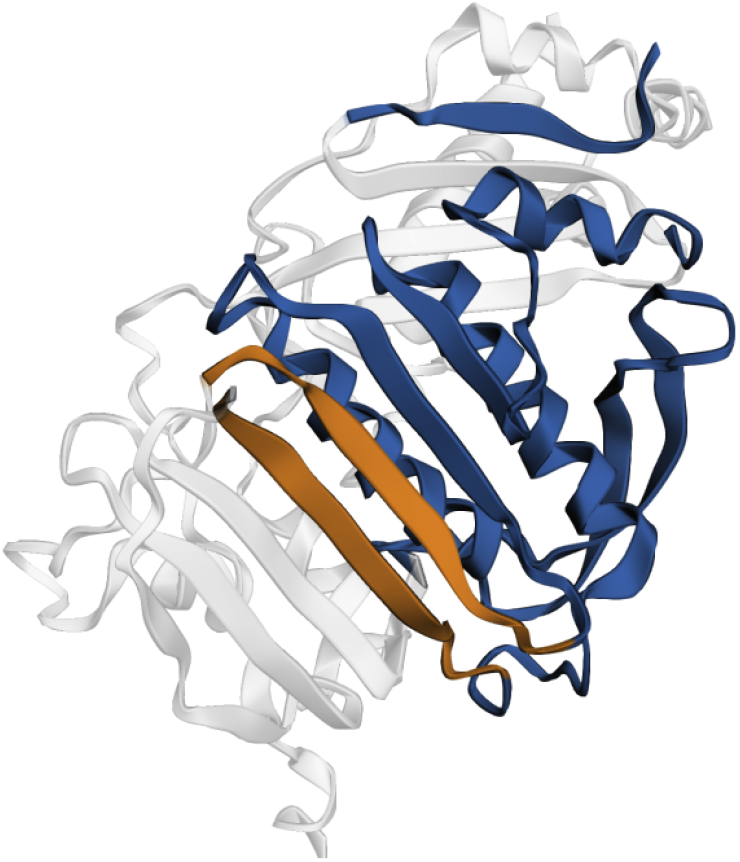
TOP2 structure (RCSB-PDB ID: 1PVG Chain A Classen et al. [2003]) with SSE elements used for contact prediction in orange, and relevant flank regions in blue [Zhang et al., 2024]

**Figure 5.**
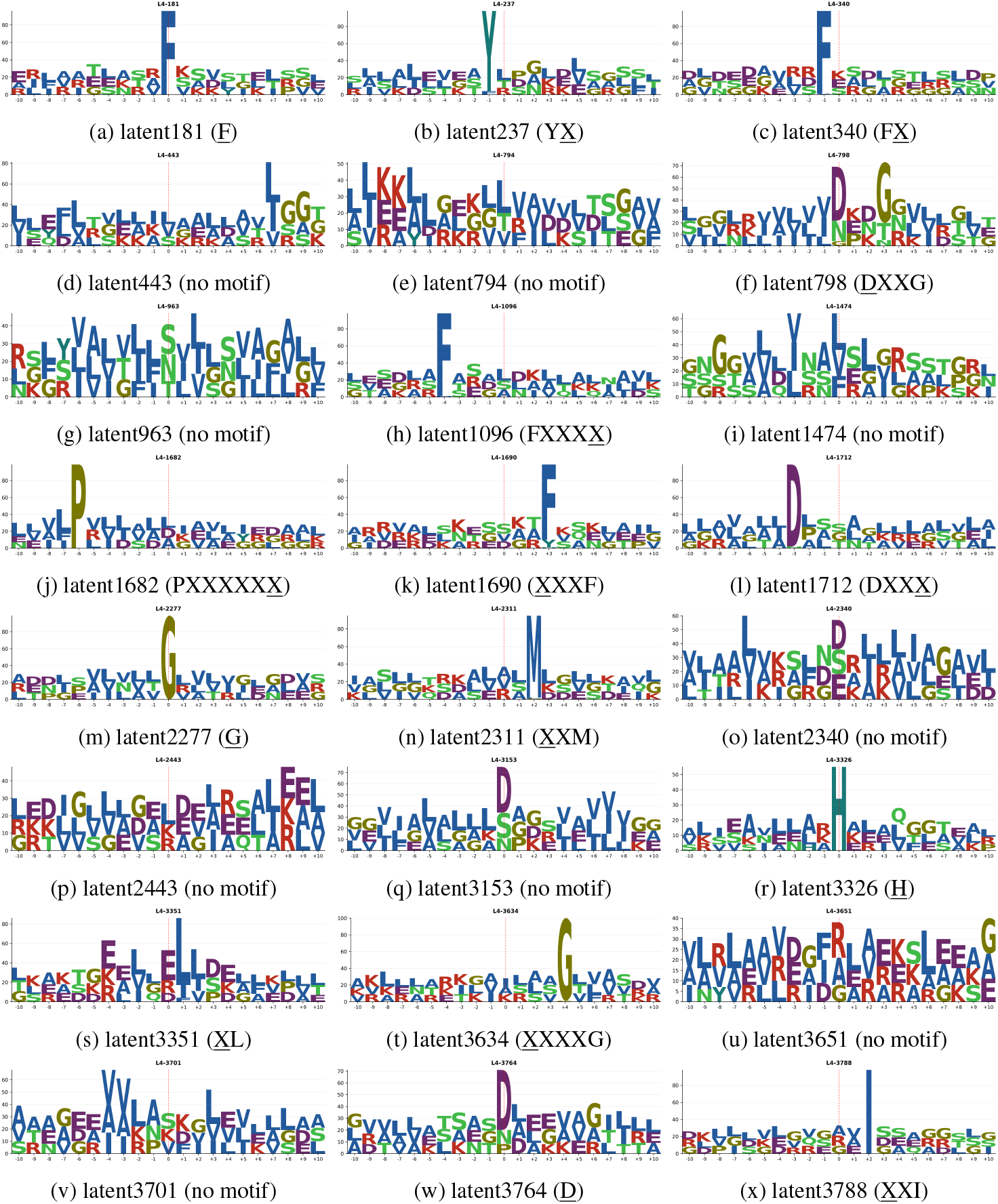
Sequence logos for motif-detector latents in layer 4 for MetXA. Each panel shows the 21-residue window centered on the max-activating token; y-axis is information (bits).

**Figure 6.**
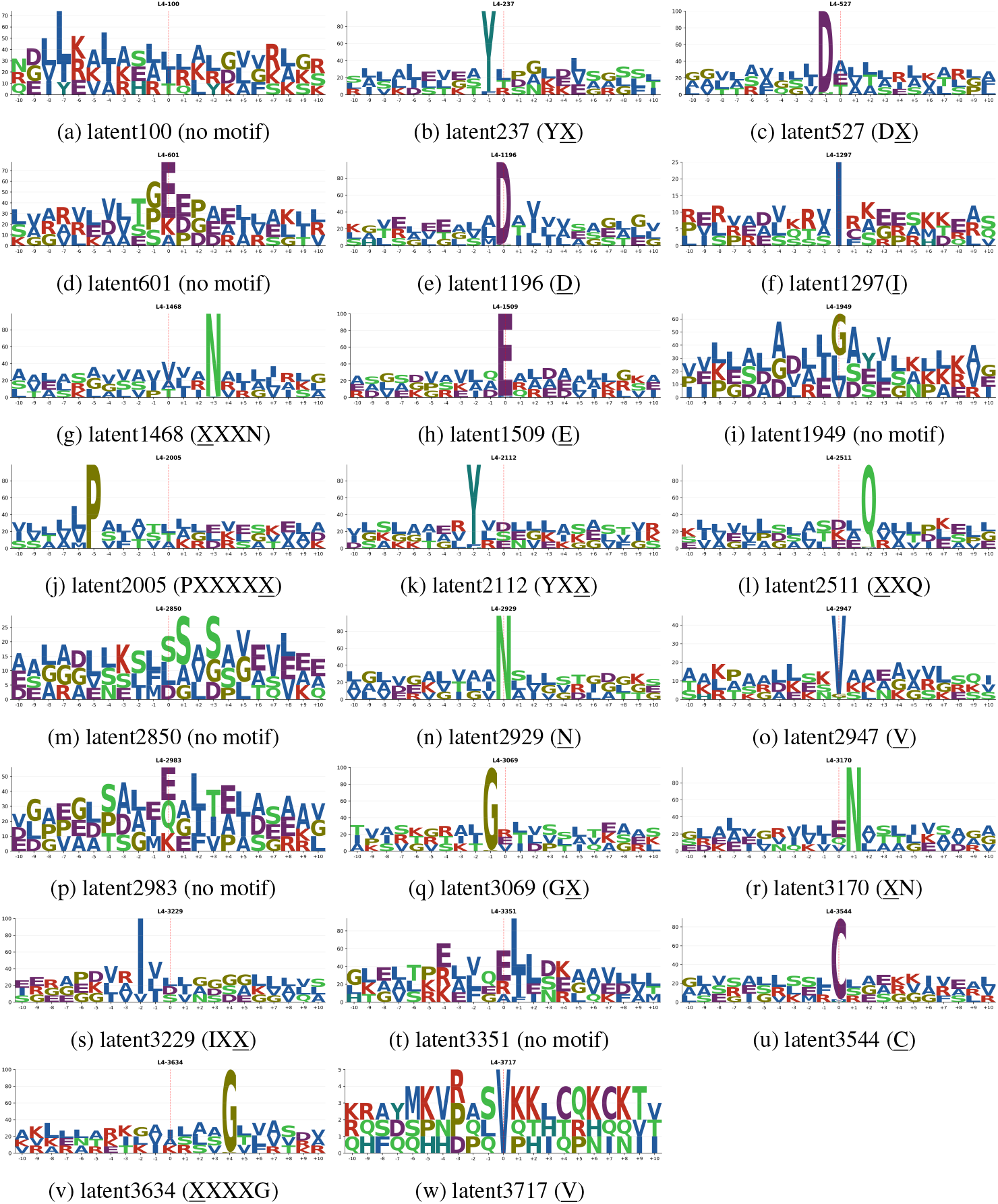
Sequence logos for motif-detector latents in layer 4 for TOP2. Each panel shows the 21-residue window centered on the max-activating token; y-axis is information (bits).

We found motif detectors not annotated during the manual analysis, including F (Layer 4, 181), FXXXX (Layer 4, 1096), and DXXX (Layer 4, 1712). The SAE latents we identified as motif detectors behave as motif detectors across the proteome, not just in our specific case study proteins.

#### 2.4.2 Domain detectors preferentially activate on assigned domains

Our case studies identified causally important latents that seem to be associated with protein domains; we tested whether 23 pre-specified latent-to-domain hypotheses (10 in MetXA, 13 in TOP2) hold on the full set of reviewed Uniprot protein entries, using length-matched negatives. Because latents activate on a per-token basis and domains are a protein-level feature, we compute a protein-level latent activation score by taking the mean of the activation on the top-*q*% (where *q*=1) of tokens for that latent. We report AUROC ± 95% stratified confidence intervals to test how selective a latent *z* is for domain *d* (Sec. 4.2.2). We found that 7/10 latent-to-domain hypotheses for MetXA and 10/13 for TOP2 had AUROC > 0.95 (95% CI width *±*0.02). In MetXA, these were latents selective for AB hydrolase fold, FAD/NAD, NAT, SAM Mtases, and WD40. For TOP2, latents with high AU-ROC were associated with Kinesin, HATPase_c, and XPG_I. A few tests returned moderate AUROC results (e.g., MetXA Transketolase AUC 0.92, DHFR 0.86, Aldolase 0.93; TOP2 two HATPase_c latents at 0.71 and 0.79), suggesting these latents may be non-specific or non-selective in some way. We note that choice of the protein-level latent activation score affects the conclusions (Figs. 7, 8,and 9). We found that the global mean dilutes signal of sparse detectors in long sequences; the max and top-K are length-biased. The top-*q* score captures signal for both sparse and dense detectors and is not length-biased.

**Figure 7.**
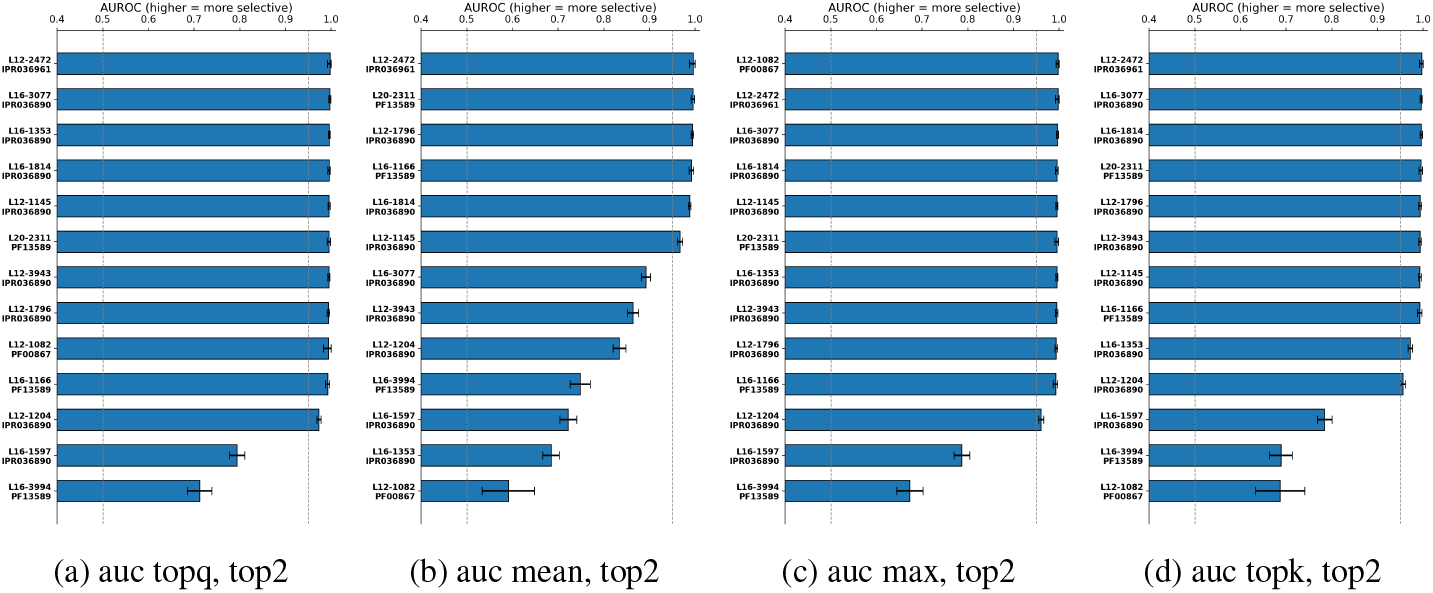
AUROC score bar charts for TOP2 using mean, max and topk aggregators.

**Figure 8.**
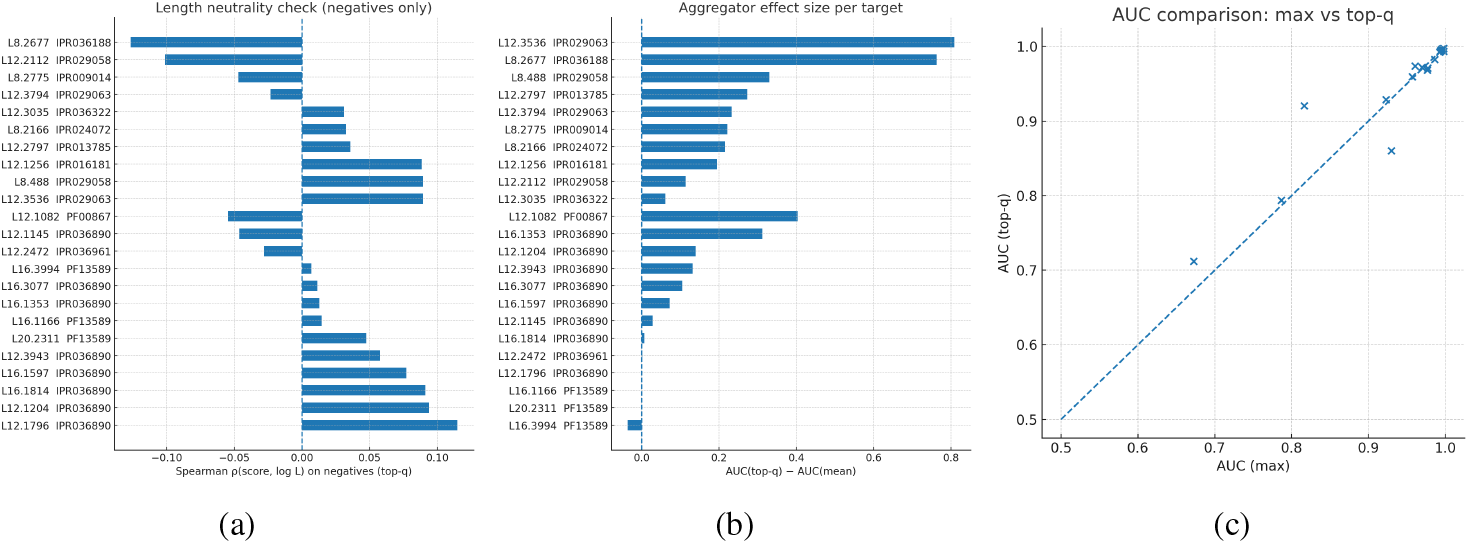
(a) Spearman rho correlation topq score vs log L, for negative samples between -0.1 to 0.1. (b) Demonstrating difference in auroc for mean and top q%. (c) AUC comparison max vs top q%.

**Figure 9.**
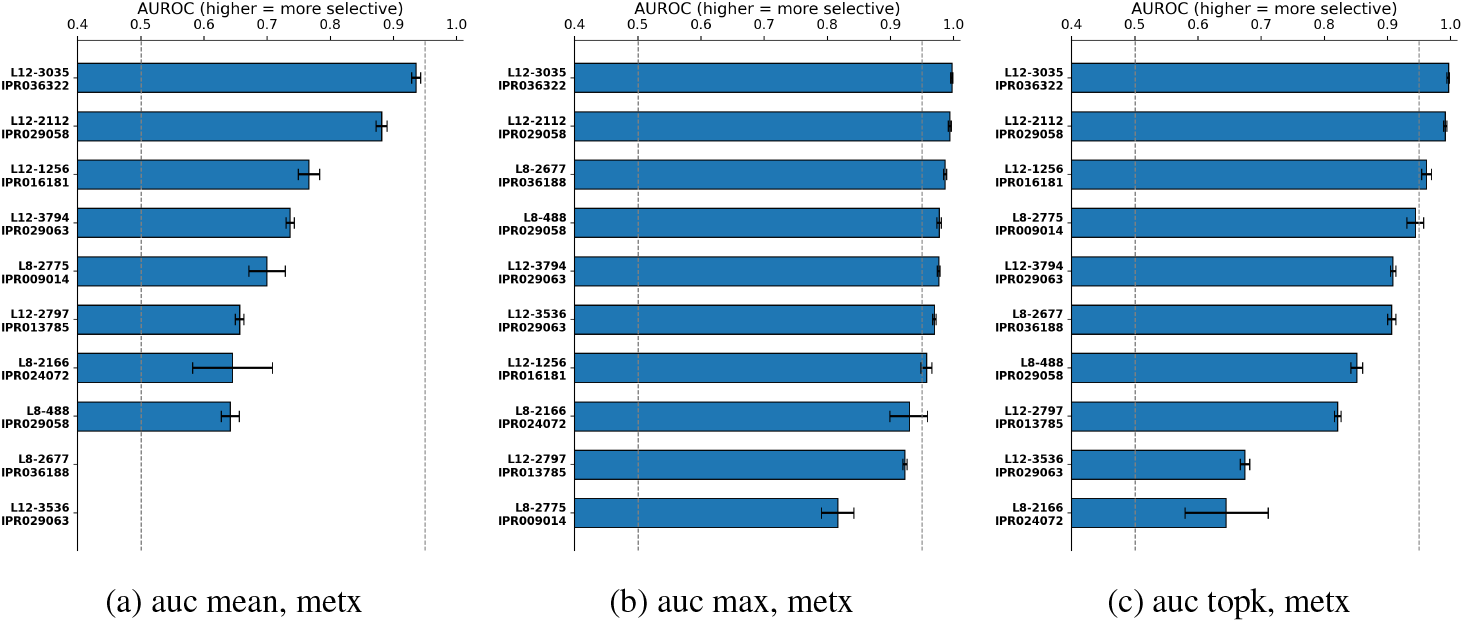
AUROC score bar charts for MetXA using mean, max and topk aggregators.

#### 2.4.3 Domain detectors are dependent on motif detectors

We hypothesized from the case studies was that domain detectors are causally gated by motif detectors. That is, the presence of specific motifs allows the model to detect the domain/family of the protein. To test this we use *path patching* [Wang et al., 2022]: *(1) patch the earlier feature to its corrupted value and record the value the later feature takes; (2) in a fresh clean run, set only that later feature to the recorded value and measure the change in contact recovery. This isolates the effect flowing along that specific link; we report edge strength as* |Δ*m*|. (Sec. B.5.3)

We rank edges by absolute effect |Δ*m*| and compute the cumulative area under the sorted curve 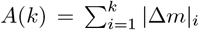 (Fig. 10, 11). We choose the smallest *k* such that *A*(*k*) ≥ 0.75 *× A*(all). Under this rule, MetXA requires *k* = 316*/*1580 edges (20.0%) to cover 76.5% AUC, and TOP2 requires *k* = 244*/*1064 edges (22.9%) to cover 75.4% AUC.

**Figure 10.**
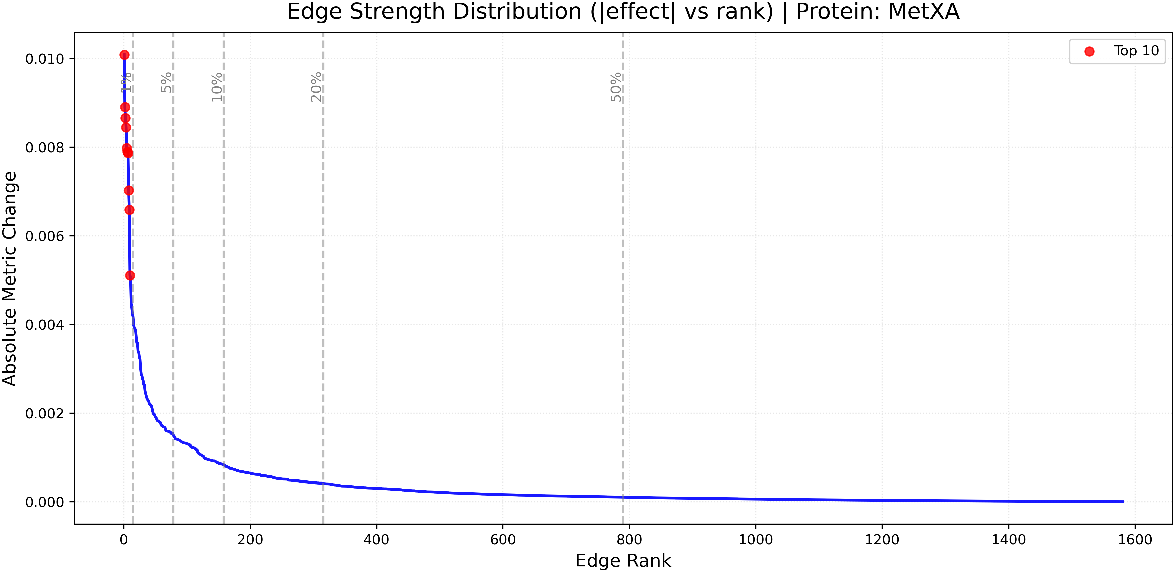
Edge strength distribution (sorted by |Δ*m*|) for MetXA.

**Figure 11.**
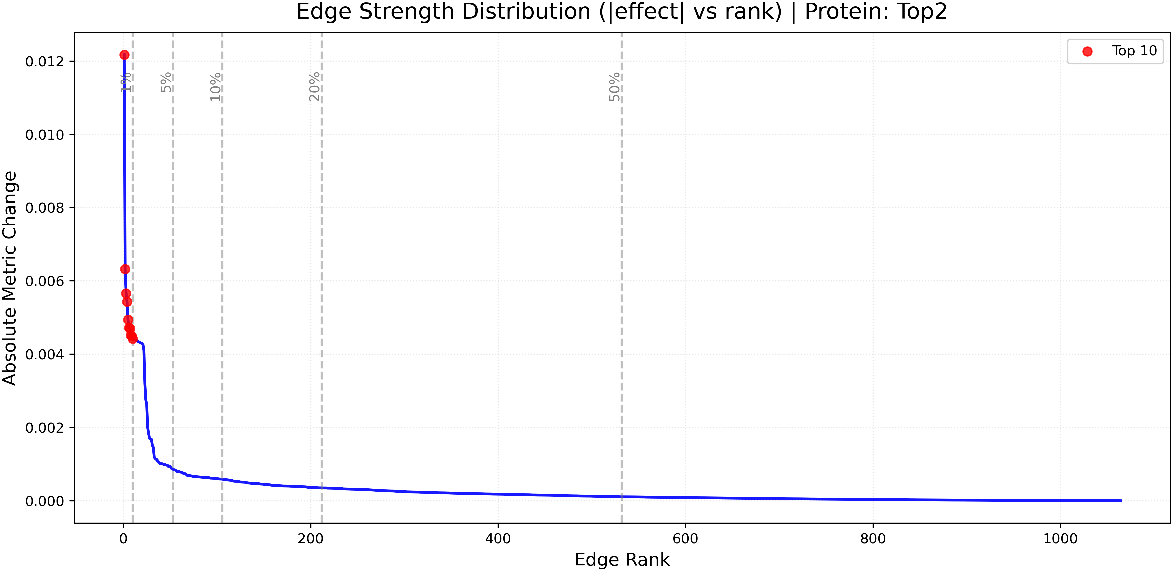
Edge strength distribution (sorted by |Δ*m*|) for TOP2.

##### Homoserine *O*-acetyltransferase

Within the AUC-75% set (316 edges; 20.0% of 1580), we find *28* edges between labeled components (Table 7), including multiple motif→domain links consistent with gating. For example, motif detectors for FX, XXI, XXM, and YX connect to AB-hydrolase detector (2112) in Layer 12 (Fig. 3)Most late-layer domain detectors receive at least one motif detector input in this set; one Layer-12 domain detector lacks a direct motif edge but connects via another Layer-12 domain that *does* receive motif input, consistent with motif-gated recognition through an intermediate domain.

**Table 7.**
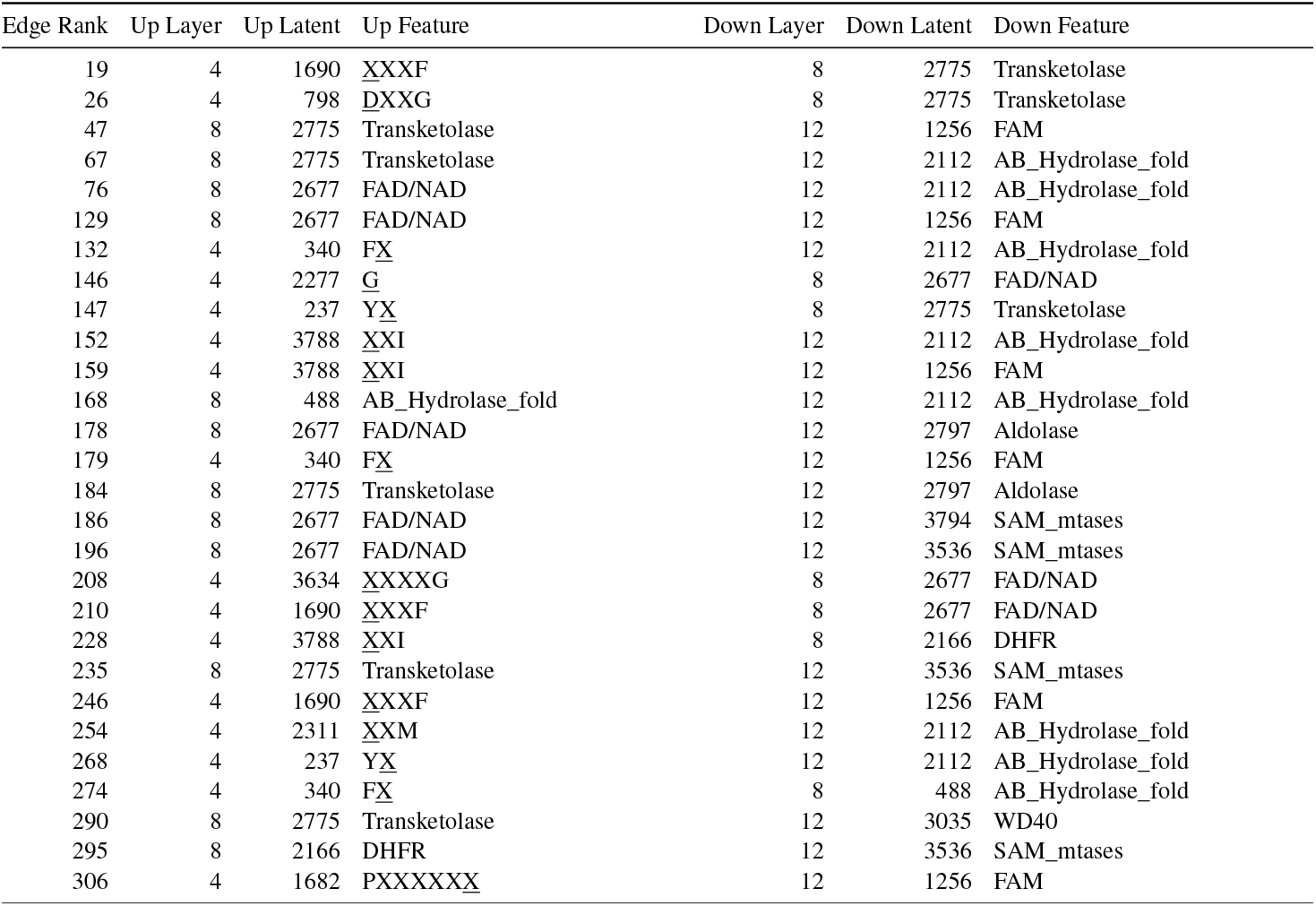
Interpretable edges for MetXA between manually analyzed latents. Upstream is earlier, downstream is later.

##### DNA topoisomerase 2

Within the AUC-75% set (244 edges; 22.9% of 1064), we observe *39* edges between labeled components (Table 8). As a representative example, several Layer-4 motif detectors converge on a *single* HATPase_c detector (1166) in Layer 16—E, XN, XXQ, V DX, XXXN, and GX—with ranks 46, 66, 79, 100, 189, 225, and 229, respectively. Other HATPase_c and Kinesin detectors in Layers 12 and 16 also receive motif detector inputs (see Table 8). All but one late-layer domain detector have a direct motif input within this set; the remaining detector has no labeled inbound edges.

**Table 8.** Interpretable edges for TOP2 between manually analyzed latents. Upstream is earlier, downstream is later.

| Edge Rank | Up Layer | Up Latent | Up Feature | Down Layer | Down Latent | Down Feature |
| --- | --- | --- | --- | --- | --- | --- |
| 1 | 12 | 3943 | Hatpase_C | 16 | 1353 | Hatpase_C |
| 25 | 12 | 1204 | Hatpase_C | 16 | 1166 | Hatpase_C |
| 28 | 12 | 1204 | Hatpase_C | 16 | 1597 | Hatpase_C |
| 32 | 12 | 1796 | Hatpase_C | 16 | 1353 | Hatpase_C |
| 34 | 4 | 1509 | E | 16 | 1353 | Hatpase_C |
| 40 | 12 | 3943 | Hatpase_C | 16 | 1597 | Hatpase_C |
| 46 | 4 | 1509 | E | 16 | 1166 | Hatpase_C |
| 48 | 12 | 1796 | Hatpase_C | 16 | 1597 | Hatpase_C |
| 54 | 4 | 1509 | E | 12 | 3943 | Hatpase_C |
| 55 | 4 | 1509 | E | 16 | 1597 | Hatpase_C |
| 57 | 4 | 3170 | XN | 12 | 3943 | Hatpase_C |
| 61 | 4 | 3170 | XN | 16 | 1597 | Hatpase_C |
| 66 | 4 | 3170 | XN | 16 | 1166 | Hatpase_C |
| 68 | 4 | 2112 | YXX | 12 | 2472 | Kinesin |
| 76 | 4 | 2511 | XXQ | 12 | 2472 | Kinesin |
| 79 | 4 | 2511 | XXQ | 16 | 1166 | Hatpase_C |
| 82 | 4 | 2929 | N | 12 | 2472 | Kinesin |
| 87 | 4 | 527 | D$\bar{X}$ | 12 | 2472 | Kinesin |
| 91 | 4 | 3717 | V | 12 | 2472 | Kinesin |
| 92 | 4 | 1468 | XXXN | 12 | 2472 | Kinesin |
| 100 | 4 | 3717 | V | 16 | 1597 | Hatpase_C |
| 102 | 4 | 3170 | XN | 12 | 2472 | Kinesin |
| 118 | 4 | 3069 | G$\bar{X}$ | 16 | 1353 | Hatpase_C |
| 123 | 4 | 3717 | V | 16 | 1166 | Hatpase_C |
| 137 | 12 | 1204 | Hatpase_C | 16 | 1353 | Hatpase_C |
| 140 | 4 | 1509 | E | 12 | 1796 | Hatpase_C |
| 151 | 12 | 2472 | Kinesin | 16 | 1597 | Hatpase_C |
| 157 | 4 | 1468 | XXXN | 12 | 3943 | Hatpase_C |
| 175 | 4 | 3170 | XN | 16 | 1353 | Hatpase_C |
| 189 | 4 | 527 | D$\bar{X}$ | 16 | 1166 | Hatpase_C |
| 198 | 4 | 3069 | G$\bar{X}$ | 12 | 1204 | Hatpase_C |
| 203 | 4 | 1468 | XXXN | 16 | 1353 | Hatpase_C |
| 204 | 4 | 1468 | XXXN | 16 | 1597 | Hatpase_C |
| 223 | 12 | 2472 | Kinesin | 16 | 1166 | Hatpase_C |
| 225 | 4 | 1468 | XXXN | 16 | 1166 | Hatpase_C |
| 229 | 4 | 3069 | G$\bar{X}$ | 16 | 1166 | Hatpase_C |
| 232 | 4 | 3544 | C | 12 | 1204 | Hatpase_C |
| 234 | 4 | 1297 | I | 16 | 1597 | Hatpase_C |
| 238 | 4 | 3544 | C | 12 | 3943 | Hatpase_C |

## 3 Discussion

We provide the first example of circuit analysis for pLMs, by adapting the causal intervention framework from mechanistic interpretability. We demonstrate how causal intervention on SAE latents using clean/corrupted input pairs can define the internal circuits used by pLMs to perform a down-stream task. While we apply our framework to contact prediction in ESM, it is readily generalizable to other pLMs and tasks. For our case study proteins, we show that preserving only a tiny fraction of latent–token pairs is sufficient to retain most post-jump accuracy. We observe a small set of early-layer latents that respond to short sequence motifs, that gate mid-to-late latents selective for protein domains/families, as confirmed by path-level ablations. We emphasize that these links are *model-internal* causal dependencies under our perturbation scheme; our work does not address *biochemical* mechanism or causality.

Identifying and labeling causally-relevant latent-token pairs could enable new forms of discovery with pLMs. First, we can check whether model predictions rely on biologically sensible evidence. Second, this will enable targeted editing and steering: attenuating misleading latents or amplifying mechanistically plausible ones, without retraining the entire model. Third, we can perform systematic follow up on cases where our labels do or do not align with known motifs and domains. This analysis may uncover overlooked functional sites, suggest previously unrecognized domain relationships, and inspire wet-lab tests that feed back into both model refinement and biological discovery.

### 3.1 Limitations

Our causal claims are restricted to *where* we intervened and *what* we measured. We study ESM-2-650M because residual-stream SAEs are publicly available for this variant, but only for every four transformer blocks. For tractability we rank and evaluate layer bottlenecks for the first 3 SAE layers, and our work is restricted to contact prediction circuits in two case-study proteins. Some top-ranked latents could not be confidently labeled as motif or domain detectors. Our motif-conservation check does not perform explicit multiple-sequence alignment and may miss gapped/shifted motifs.

#### 3.2 Future Work

Future work will expand the scope of our causal annotation for contact prediction. SAEs trained at every layer (and ideally on attention/MLP streams) or cross layer transcoders [Lindsey et al., 2025], would enable cross-layer minimal-set searches for the full circuit. Extending our analysis from the flank-induced jumps to full input sequences will test whether the same motif → domain logic persists when many residue pairs are jointly scored, and whether additional long-range features emerge. Finally, rather than summarizing interventions with a single scalar, we will analyze *per-contact* effects: which residue pairs gain/lose probability under targeted latent edits, how these changes cluster in 2D contact space (e.g., within/between SSEs), and how they project to 3D via structure prediction.

We plan automated labeling of latent-token pairs to reduce manual effort and improve label reliability. We will then seek to generalize across proteins and scales, to reveal which motif/domain detectors and dependencies are shared vs. protein-specific, how they shift with model size, and whether “domain-labeled” late-layer units sometimes act as short-motif proxies.

Together, these directions take us from a tractable layer-wise bottleneck to a complete, cross-layer circuit for contact prediction, and from two case studies toward a library of mechanistic explanations that are auditable, reusable, and biologically informative.

## 4 Methods

In this section, we cover the methods used for case study analysis §2.3 and selectivity tests §2.4.1,2.4.2. We overview the model and data selection in Appendix §B.1, contact prediction task in Appendix §B.2, the causal intervention framework in Appendix §B.5.

### 4.1 Latent Interpretation and Case-Study Analysis

Each latent–token pair in the layer-wise bottlenecks was manually examined for two complementary properties: its global role and its jump-specific role. For the global role, we select the 20 UniRef50 proteins with the highest activation for the latent, and load them in the InterProt viewer [InterProt Team, 2025] [access date: June 30, 2025] annotations that overlap with the token position were retrieved automatically from UniProt and recurring sequence motifs were noted by eye. For the jump-specific role, we plotted the latent’s per-token activations on the case-study protein under three inputs (corrupted, clean, and fully unmasked). The indirect-effect ranking already scales with activation change, so no additional numeric threshold was imposed. Latents whose motif encompassed a residue newly revealed in the clean input were labeled directly explained motif detectors; those whose motif lay elsewhere were labeled indirect motif detectors. Finally, latents whose activation patterns matched a specific InterPro domain—regardless of whether that domain is annotated for the target protein—were labeled domain detectors.

### 4.2 Quantitative latent labeling

From the full set of UniProt reviewed proteins (*n* = 573,661 as of June 30, 2025), we first filtered out any proteins with sequence length *>* 1022, as ESM-2 adds two extra tokens to the input and caps input length at 1024 tokens. Then, we randomly selected a set of 10,000 proteins from this length-restricted subset. For each of these proteins and for each available latent, the per-token activation was extracted and stored. Using the per-token activations, we then quantitatively labeled latents for both motifs and domains.

#### 4.2.1 Motif Labeling through Sequence Conservation

To identify sequence motifs associated with latent activations, we aligned sequences at their maximum activation positions and analyzed conservation patterns in flanking regions. For each latent, we identified the highest-activating residue in each protein, ranked these maxima across the dataset, selected the top 100 residues (ensuring each protein was represented only once), and computed position-specific conservation scores to quantitatively characterize activation-associated motifs. Specifically, for each of those top 100 residues, we extract the 10 amino acids before and after it in the sequence, truncating if we run into the beginning or end of the protein. The window size was chosen empirically based on manual inspection. For each latent, we then created a sequence logo using those 100 sequences of length 21 (10+1+10). Sequence logos were generated with the LOGOMAKER Python library [Tareen and Kinney, 2020]. The *x*-axis of such a logo gives the sequence position, relative to the middle residue (the highest-activating residue). The *y*-axis of a sequence logo gives the information content in bits, where the height *h* of an amino acid *a* at position *i* is given by:

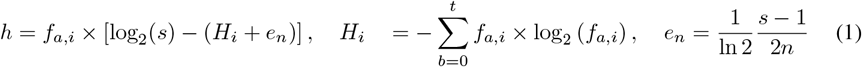

where *f*_*a,i*_ is the relative frequency of amino acid *a* at position *i* and *s* = 20 because we are only considering the 20 canonical amino acids. The quantity *H*_*i*_ is the Shannon entropy or the uncertainty of position *i*, and *e*_*n*_ is the small-sample correction for an alignment of *n* letters (here, *n* = 100 sequences) [Schneider and Stephens, 1990]. The final sequence logos for each latent represent a statistical view of the conservation of amino acids at each position in the neighborhood of the highest-activated residue.

#### 4.2.2 Domain labeling through correlation

##### Hypothesis-driven

Protein domains are conserved sequences associated with structure and function [Chothia, 1992], and are curated in InterPro [Blum et al., 2025]. For each latent *z* and domain *d*, we hypothesize score(*X, z*) is higher for proteins (*X*) with *d* than without *d*. We test these pre-specified latent-to-domain hypotheses on a shared dataset. For a domain *d*, positive examples are randomly selected from the set of UniProt reviewed proteins with domain *d*. We defined a candidate negative pool as reviewed Uniprot proteins without *d*, and length-matched negatives to positives via stratified sampling over 12 empirical length-quantile bins to remove sequence-length confounding. Quantiles are computed on the union of positive and candidate-negative lengths for that domain. Within each bin, we sampled negatives without replacement up to the number of positives in that bin; if fewer negatives were available, we took all available and accepted minor imbalance. We summarize a protein’s activation score by the *mean of the top q%* of tokens, with *q* = 1 (fixed *a priori*). This captures both sparse and dense signals without being length biased. An ideal score metric should also expect low Spearman Rank Correlation rho (*ρ*(score, log*L*) ≈ 0) on negatives. This is because for proteins without domain *d*, a good per-protein score should not systematically increase/decrease with sequence length. With this setup, we calculated the AUROC using scikit-learn [Pedregosa et al., 2011] to measure the effect strength for selectivity and provide 95% stratified bootstrap confidence intervals (CIs). AUROC denotes the probability that a random positive out-scores a random negative; CIs use 3000 stratified bootstrap resamples. For comparison, we computed the same metrics for max, mean, top-K as for top-*q*. We found that the global mean dilutes sparse activations, while the max and top-K had higher length correlations for negatives (Appendix §D).

## Acknowledgments and Disclosure of Funding

The authors thank members of the UMass SAGE lab for valuable input. This work utilized resources from Unity, a collaborative, multi-institutional high-performance computing cluster managed by UMass Amherst Research Computing and Data. BMR was supported by the UMass Spaulding-Smith Fellowship.

## 4 DNA topoisomerase 2 Layer wise analysis

### Layer 4

The zero-circuit performance is *m*_0_ = 0.33; thus the explainable window is *W* = *T ™ m*_0_ = 0.27. The circuit requires 29 latent–token pairs to meet the criterion. Similar to the results for MetXA, observe motif detectors. This cluster contains 13 pairs (44.8% of the layer). *Direct* motif detectors (20.7% of layer). Ablating them reduces *m*(*X*) by 6.3% of *W* (0.017). Example: a latent at G34 fires on YXX across its top-20 proteins (E1) and deactivates when Y28 is masked (E2→Q2). *Indirect* motif detectors (24.1% of layer). Ablating them reduces *m*(*X*) by 11% of *W* (0.03). Example: a latent at I101 prefers IXX (E1); its activation rises only in the clean input (E2→Q2). Together, motif detectors accounts for 19.10% of *W* (0.053).

### Layer 12

The zero-circuit performance is *m*_0_ = 0.096; thus the explainable window is *W* = *T ™ m*_0_ = 0.50. The circuit requires 18 latent–token pairs to meet the criterion. This layer contains *domain detectors*. This cluster contains 5 pairs (27.8% of the layer). *Annotated* domains (TOP2 ‘s own labels: GHKL / HATPase_c) (16.7% of layer). Ablating them reduces *m*(*X*) by 24.6% of *W* (0.125).

*Misc* domains (e.g., XPG-I or Kinesin) (11.1% of layer). Ablating them reduces *m*(*X*) by 1.6% of *W* (0.01). Example: two latents prefer XPG-I/Kinesin families (E1); removal yields a small drop. Together, domain detectors accounts for 31.3% of *W* (0.158).

### Layer 16

The zero-circuit performance is *m*_0_ = 0.11; thus the explainable window is *W* = *T ™ m*_0_ = 0.49. The circuit requires 25 latent–token pairs to meet the criterion. This layer also contains *domain detectors*. This cluster contains 6 pairs (24% of the layer). *Annotated* (GHKL / HATPase_c) (24% of layer). Ablating them reduces *m*(*X*) by 15.6% of *W* (0.0785). Example: a latent at V201 fires on GHKL proteins (E1) and activates only in the clean run (E2→Q2). Together, domain detectors accounts for 15.6% of *W* (0.0785).

## Materials

### Model and Data Selection

#### Protein language model

All experiments use ESM-2-650M as the primary protein language model, from FAIR’s public repository (“esm2_t33_650M_UR50D” checkpoint). No additional fine-tuning was performed [Lin et al., 2023b, Meta AI (FAIR), 2023b].

#### Sparse autoencoders (SAEs)

We use eight publicly released SAEs from Adams et al. [2025], trained on residual stream activations from layers 4, 8, 12, 16, 20, 24, 28, and 32. Each SAE encodes the 1 280-dimensional residual activation (extracted after the attention and MLP sublayers) to a 4 096-d latent vector *z*, followed by a TopK gate (*k* = 64) to enforce sparsity. Throughout, *latents* refers exclusively to these pretrained SAE features *z*.

#### Proteome for selectivity assays

AUROC and enrichment tests are applied to the reviewed *Swiss-Prot* subset of UniProt (*N* = 573, 661 proteins) [Boutet et al., 2007]. For manual inspection of top-activating sequences we use the InterProt viewer (UR50 protein set, accessed 2 Aug 2025).

#### Case-study proteins

Zhang et al. [2024] demonstrated the sudden increase in contact recovery on ESM-2-3B. As both open sources SAEs were only available on ESM-2-650M, we iterated over the set of proteins from Zhang et al. [2024] and picked the two that showcased the jump in the smaller 650M model. (1) *DNA topoisomerase 2*; Species: *Saccharomyces cerevisiae strain ATCC 204508 / S288c (baker’s yeast)* S288C; UniProt: P06786; PDB: 1PVG [Mirza et al., 2005] (2) *Homoserine O-acetyltransferase*; Species: *Haemophilus influenzae strain ATCC 51907 / DSM 11121 / KW20 / Rd*; UniProt: P45131; PDB: 2B61 [Classen et al., 2003].

### B.2 Contact-prediction task

We adapt the contact prediction task defined by Zhang et al. [2024], using the ESM-2 contact prediction head [Rao et al., 2020, Meta AI (FAIR), 2023a]. They built a dataset of pairs of contacting secondary-structure elements (SSEs) separated by *>* 100 aa in the primary sequence across 266 proteins. For an input protein sequence *X* (which may contain masked tokens) and its corresponding pair of SSEs from their database, we define the per-SSE-pair prediction quality as the *recovery score*

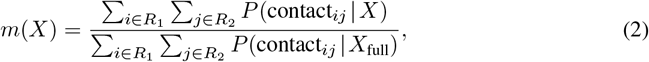

where *R*_1_, *R*_2_ index the two SSEs and *X*_full_ is the unmasked sequence.

#### Jump setup

Two inputs are compared: a *clean* sequence whose flanks yield high recovery (typically, *m ≈* 1) and a *corrupted* sequence with two fewer flank residues that collapses to near-random (*m* ≲ 0.1). The step-change constitutes is referred to as the jump.

### B.3 Motif notation

We describe motifs using one-letter amino acid codes with the following notation:

- {A,C,D,E,F,G,H,I,K,L,M,N,P,Q,R,S,T,V,W,Y}: a specific, conserved amino acid.
- X: any amino acid (position not conserved).
- <u>underline</u>: the token at which the latent activates.

### B.4 Sparse Autoencoders (SAEs)

Neurons in deep neural networks and language models are usually polysemantic, meaning that they activate on to multiple unrelated variables or concepts [Olah et al., 2020]. One potential cause of polysemanticity is superposition, where a neural network represents more independent features of the data than it has neurons for by assigning each feature its own linear combination of neurons [Bricken et al., 2023]. The work by Elhage et al. [2022] has shown how Sparse Autoencoders (SAEs) can be used to disentangle these dense representations into monosemantic neurons, which represent a single concept/variable.

We use the sparse autoencoders trained by Adams et al., which followed Gao et al., where each SAE is a linear encoder–decoder that learns a sparse, length-*k* latent vector *z* for every residual-stream activation *x* ∈ ℝ^*d*^:

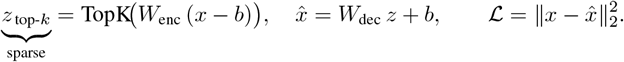

Individual neurons *z*_*i*_ in the SAE are referred to as latents. Because only the *k* top activating latents survive the TopK gate, individual latents are often monosemantic [Gao et al., 2024], making them easier to interpret or ablate.

### B.5 Circuit Discovery

#### B.5.1 Causal influence ranking by activation patching

We aim to discover which specific latents at each sequence position (latent–token pairs) are causally necessary for the contact prediction jump. First, we conduct two forward passes of the network, one with the clean sequence that produces near-complete contact recovery, and one with the corrupted sequence that produces near-zero contact recovery. Then, for each SAE latent at each sequence position, we measure its *indirect effect (IE)* [Pearl, 2022*] using counterfactual activation patching [Vig et al., 2020, Finlayson et al., 2021]*. *Activation patching copies the hidden activations of a single network component from one forward pass into another. Here, we copy the latent activations from the corrupted forward pass into a clean forward pass. The change in the model’s output that results from patching a component is called the indirect effect (IE)* of that component.

##### Indirect-effect calculation

For a component **a** we patch its activations from the failing run into the successful one and recompute the score: *m* (*X*_clean_ | do(**a** ← **a**_patch_)).

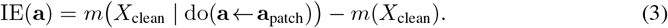

Patching can be restricted to a single position *t* by swapping only **a**[*t*]. We refer to the activation of a latent at a specific sequence position as “latent–token pair.” Components are ranked by the magnitude of their IE; a large *negative* value indicates that importing **a** from the corrupted run alone has a large negative impact on contact recovery even though the rest of the network still receives the clean input.

Directly evaluating the indirect effect in (3) for *every* latent–token pair would require a forward pass per component (*∼* 10^7^), which is infeasible. Instead, following Marks et al. [2024] we compute effects with two gradient-based approximations: **Attribution Patching** [Nanda, 2023, Syed et al., 2023, Kramár et al., 2024]: a first-order Taylor expansion around the clean run that estimates the effect of *all* components using only two forward passes and a single backward pass. **Integrated Gradients** [Sundararajan et al., 2017, Hanna et al., 2024]: a more accurate path integral of the gradient along the straight-line interpolation between patched and clean activations. We use *N* =10 [Marks et al., 2024] evenly spaced interpolation points, trading the extra 10 forward–backward pairs for a noticeably tighter fit than only using attribution patching Hanna et al. [2024].

#### B.5.2 Circuit Discovery

Circuits are a subgraph of a neural network [Olah et al., 2020, Wang et al., 2022]. In the context of this work, we define circuit as the minimal set of latent–token pairs needed to maintain a threshold of the contact recovery, while all other pairs are ablated (frozen to their corrupted activations). We rank 4,092 *×* 8 latents at each of the *∼*400 sequence positions for IE, yielding *∼*1.3 *×* 10^7^ latent–token pairs. However, SAEs are usually not able to reconstruct the activations with 100% accuracy. So error terms [Marks et al., 2024] are added to the reconstructions to maintain performance. Mathematically, the error term *ε*(*x*) is the difference between the model and reconstructed activations:

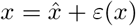

We construct the circuit with the top *K* latent–token pairs with highest indirect effect. We begin by patching the corrupted activations for every pair not in the top *K*. Then, we allow the top *K* to be recomputed during the forward pass. If contact recovery jumps, the retained pairs are sufficient to create the circuit; if it remains low, the circuit is still missing critical pieces and we continue adding more clean activations. We set the threshold at 70% of the post-jump recovery. We chose 70% to explain the majority of the jump while focusing on the most important latents.

##### Layer-wise bottlenecks

In addition to the model-spanning circuit defined above, we seek to compute layer-specific bottlenecks, defined as the minimal set of latent–token pairs from a specific layer needed to maintain a threshold of contact recovery, with all other latent–token pairs from that layer ablated and with all other layers not directly intervened. For layer *ℓ* we allow only its top-*K*_*ℓ*_ latent– token pairs to be recomputed and patch the corrupted activation for all other latent–token pairs in that layer. All other layers are also allowed to be recomputed. Caples et al. [2025]. *K*_*ℓ*_ is the smallest value reaching ≥70% recovery, this set of pairs are referred to as ℬ_*ℓ*_. Layer bottlenecks trade completeness for interpretability: ℬ_*ℓ*_ ⊆ circuit, but each is small enough for manual inspection.

#### B.5.3 Path Patching

To quantify how strongly an *upstream* SAE feature *u*_*i*_ in layer *ℓ* influences a *downstream* feature *d*_*j*_ in layer *ℓ*^*′*^, we follow Wang et al. [2022] and compute an *edge attribution* via *path patching*. This method isolates the causal pathway from *u*_*i*_ to *d*_*j*_ through a two-stage intervention:

1. **Record downstream change**: Patch the upstream feature activation from 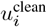 to 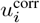 while keeping all other upstream features at their clean values. Record the resulting downstream activation 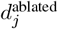.
2. **Isolate pathway effect**: In a fresh forward pass on the clean input, patch only the downstream feature from 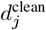 to 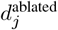 (the value recorded in step 1), keeping all other features at their clean values.
3. **Measure metric change**: Compute the change in metric (2): *w*_*i*→*j*_ = *m*_patched_ − *m*_clean_.

The resulting edge weight *w*_*i→j*_ isolates the causal contribution of feature *u*_*i*_ to the model’s performance that flows specifically through feature *d*_*j*_, excluding any parallel pathways. This twostage patching procedure ensures we capture only the direct *u*_*i*_ *→ d*_*j*_ influence, providing a precise measure of feature interaction strength. However, this needs 𝒪(*K*^2^) forward passes for complete pairwise analysis.

## C Sequence Logos

## D Domain Correlation Tables

## E Path Patching results

## F Additional recovery curves

**Figure 13.**
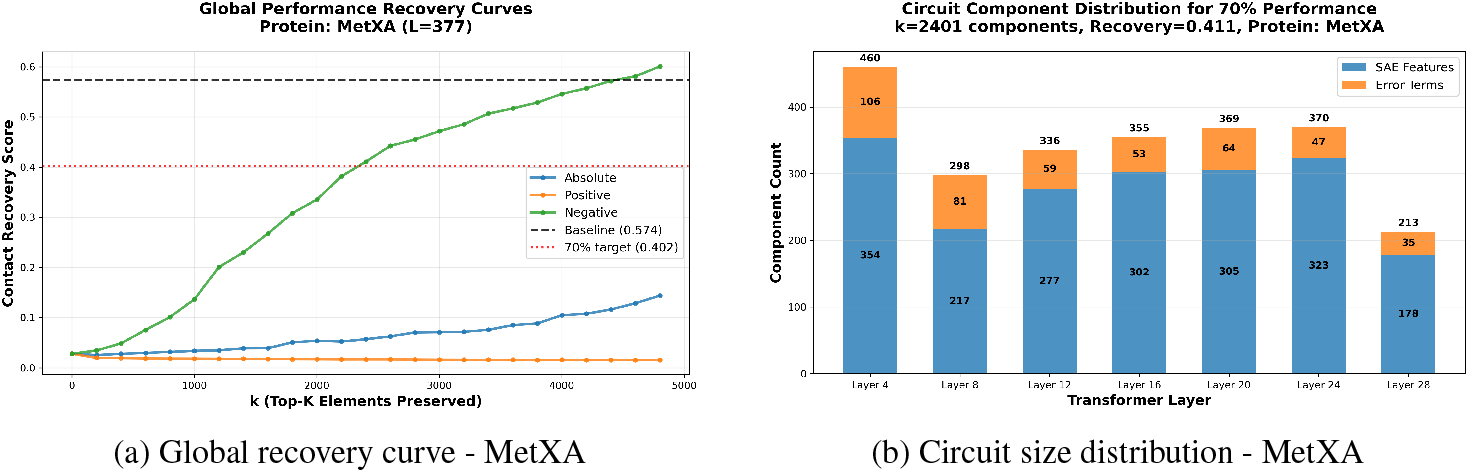
Performance and distribution diagnostics for the learned circuit.

**Figure 14.**
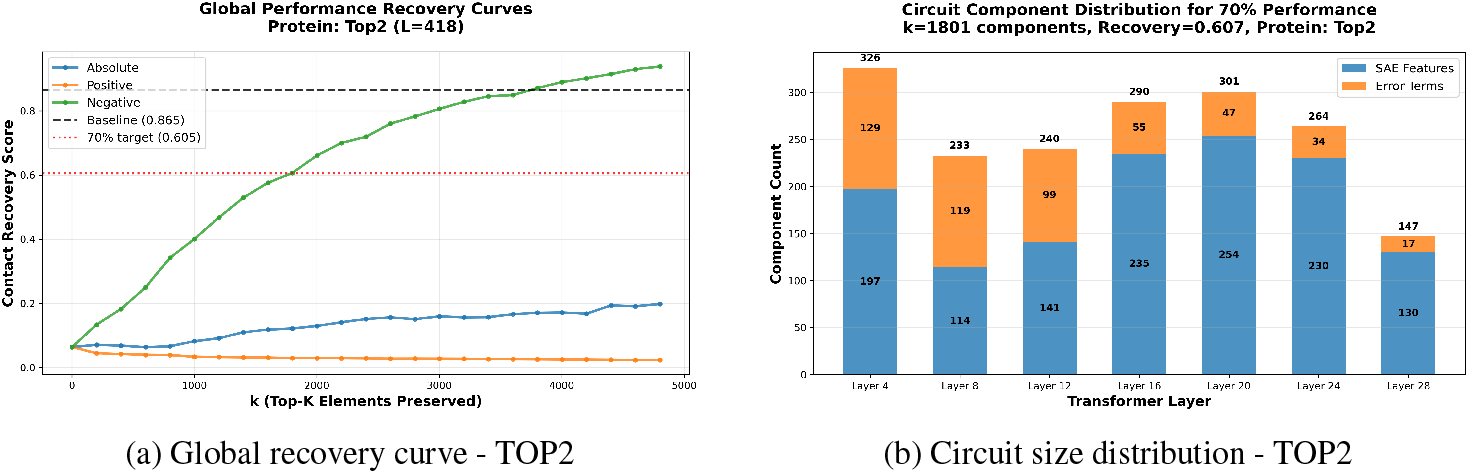
Performance and distribution diagnostics for the learned circuit.

